# Myocardial deformation imaging by 2D speckle tracking echocardiography for assessment of diastolic dysfunction in murine cardiopathology

**DOI:** 10.1101/2022.08.06.503061

**Authors:** Lorna J. Daniels, Calum Macindoe, Parisa Koutsifeli, Marco Annandale, Antonia J. A. Raaijmakers, Kate L Weeks, James R. Bell, Johannes V. Janssens, Claire L. Curl, Lea M.D. Delbridge, Kimberley M. Mellor

**Affiliations:** Department of Physiology, University of Auckland, New Zealand; Department of Anatomy and Physiology, University of Melbourne, Australia; Auckland Bioengineering Institute, University of Auckland, New Zealand; Radcliffe Department of Medicine, OCDEM, University of Oxford, United Kingdom; Baker Department of Cardiometabolic Health, University of Melbourne, Australia; Department of Microbiology, Anatomy, Physiology & Pharmacology, La Trobe University, Australia

**Author notes:** **Corresponding Author:** Kimberley M. Mellor, Cellular and Molecular Cardiology Laboratory, Department of Physiology, University of Auckland, New Zealand.

**Keywords:** speckle tracking echocardiography, myocardial wall deformation, strain, diabetes, diastolic dysfunction

## Abstract

Diastolic dysfunction is increasingly identified as a key, early onset subclinical condition characterizing cardiopathologies of rising prevalence, including diabetic heart disease and heart failure with preserved ejection fraction (HFpEF). Diastolic dysfunction characterization has important prognostic value in management of disease outcomes. Validated tools for *in vivo* monitoring of diastolic function in rodent models of diabetes are required for progress in pre-clinical cardiology studies. 2D speckle tracking echocardiography has emerged as a powerful tool for evaluating cardiac wall deformation throughout the cardiac cycle. The aim of this study was to examine the applicability of 2D speckle tracking echocardiography for comprehensive global and regional assessment of diastolic function in a pre-clinical murine model of cardio-metabolic disease. Type 2 diabetes (T2D) was induced in C57Bl/6 male mice using a high fat high sugar dietary intervention for 20 weeks. Significant impairment in left ventricle peak diastolic strain rate was evident in longitudinal, radial and circumferential planes in T2D mice. Peak diastolic velocity was similarly impaired in the longitudinal and radial planes. Regional analysis of longitudinal peak diastolic strain rate revealed that the anterior free left ventricular wall is particularly susceptible to T2D-induced diastolic dysfunction. These findings provide a significant advance on characterization of diastolic dysfunction in a pre-clinical mouse model of cardiopathology and offer a comprehensive suite of benchmark values for future pre-clinical cardiology studies.

## Introduction

Diastolic dysfunction is characterized by impaired heart relaxation and subsequent reduced ventricular filling due to increased myocardial stiffness. Identification of early diastolic dysfunction has important prognostic value for progression to heart failure. Heart failure with preserved ejection fraction (HFpEF), a condition characterized by diastolic dysfunction, is highly prevalent in diabetic patients and has worse prognosis than in non-diabetic patients^1,2^. Currently there are no effective treatments to specifically target diastolic dysfunction, thus a substantial research effort is underway for drug-discovery in pre-clinical models. There is an urgent need for validated tools for systematic evaluation of diastolic function in small rodent models of disease. Conventional echocardiography approaches to assess diastolic function clinically (e.g. flow and tissue Doppler imaging, E/e’) have been applied to tracking diastolic function in small rodents^3,4^, but high heart rates (>400 bpm) and small cardiac anatomy present unique challenges^5^. A high level of training and expertise is required to generate reliable outcomes. Speckle tracking echocardiography is a post-processing tool to track deformation of the myocardial wall throughout the cardiac cycle, and is increasingly used in both clinical^6-9^ and pre-clinical studies^10-13^. Global cardiac strain is the most commonly reported variable from speckle tracking studies, which is typically associated with characteristics of systolic function. Notably, speckle tracking-derived strain rate can be determined over the full length of the cardiac cycle and peak strain rate during the diastolic period may provide a robust additional measure to support the evaluation of diastolic function in small rodent models for pre-clinical studies^5^.

Speckle tracking echocardiography is based on tracking the motion of speckle patterns created by interference of ultrasound beams in the myocardium over time^14^. The extent and speed of deformation of a specific area of the myocardium is recorded, in relation to the initial dimensions. A significant advantage of speckle tracking echocardiography is that cardiac function can be assessed in long and short axis viewing planes to obtain information on longitudinal, radial and circumferential strain for any given area of interest. Tracking regional changes in myocardial wall deformation has proven to be useful particularly in identifying ischemic regions in studies of myocardial infarction^12,13^. This approach may also be applicable to generating a more advanced understanding of regional cardiac dysfunction in chronic disease states.

Whilst speckle tracking echocardiography is becoming more widely used, particularly for reporting global strain measures, its utility in expanding the available tools for assessment of diastolic function in small rodent models has not been fully realized. Given the escalating prevalence of diabetes, and the predisposition for diastolic dysfunction (and HFpEF) in diabetic patients, diastolic strain rate may be an important measure for pre-clinical studies investigating novel drug targets for treatment of diabetic heart disease. The few studies available reporting diastolic strain rate in diabetic rodent models have mostly used genetic models of advanced diabetes^15-17^. Regional differences in diastolic function have not been examined. An evaluation of speckle tracking echocardiography to monitor diastolic function in more clinically relevant models of diabetes is lacking. Thus the goals of this study were to validate speckle tracking echocardiography as a tool to assess diastolic myocardial wall deformation and evaluate regional differences in diastolic dysfunction in a dietary (high fat high sugar) mouse model of obesity and type 2 diabetes (T2D).

## Results

### Doppler echocardiography confirms diastolic dysfunction in T2D mice

To develop a mouse model of T2D, mice were fed a high fat high sugar diet for 20 weeks. T2D mice exhibited significantly increased bodyweight from 2 weeks of dietary intervention (Fig. 1A), impaired glucose tolerance (11 weeks diet duration, Fig. 1B), and hyperglycemia at study endpoint (20 weeks diet duration, Fig. 1C). Representative pulse wave and tissue Doppler images from control and T2D mice are presented in Fig. 1D. T2D mice showed no change in the ratio of early (E) to late (A) mitral valve blood flow rate (E/A ratio) compared to control mice (Fig. 1E). The commonly used clinical index of ventricular filling (ratio of early (E) mitral valve flow rate to early mitral annulus tissue movement (e’), E/e’) was significantly increased (Fig. 1F), confirming the presence of diastolic dysfunction in this dietary T2D mouse model.

**Figure 1.**
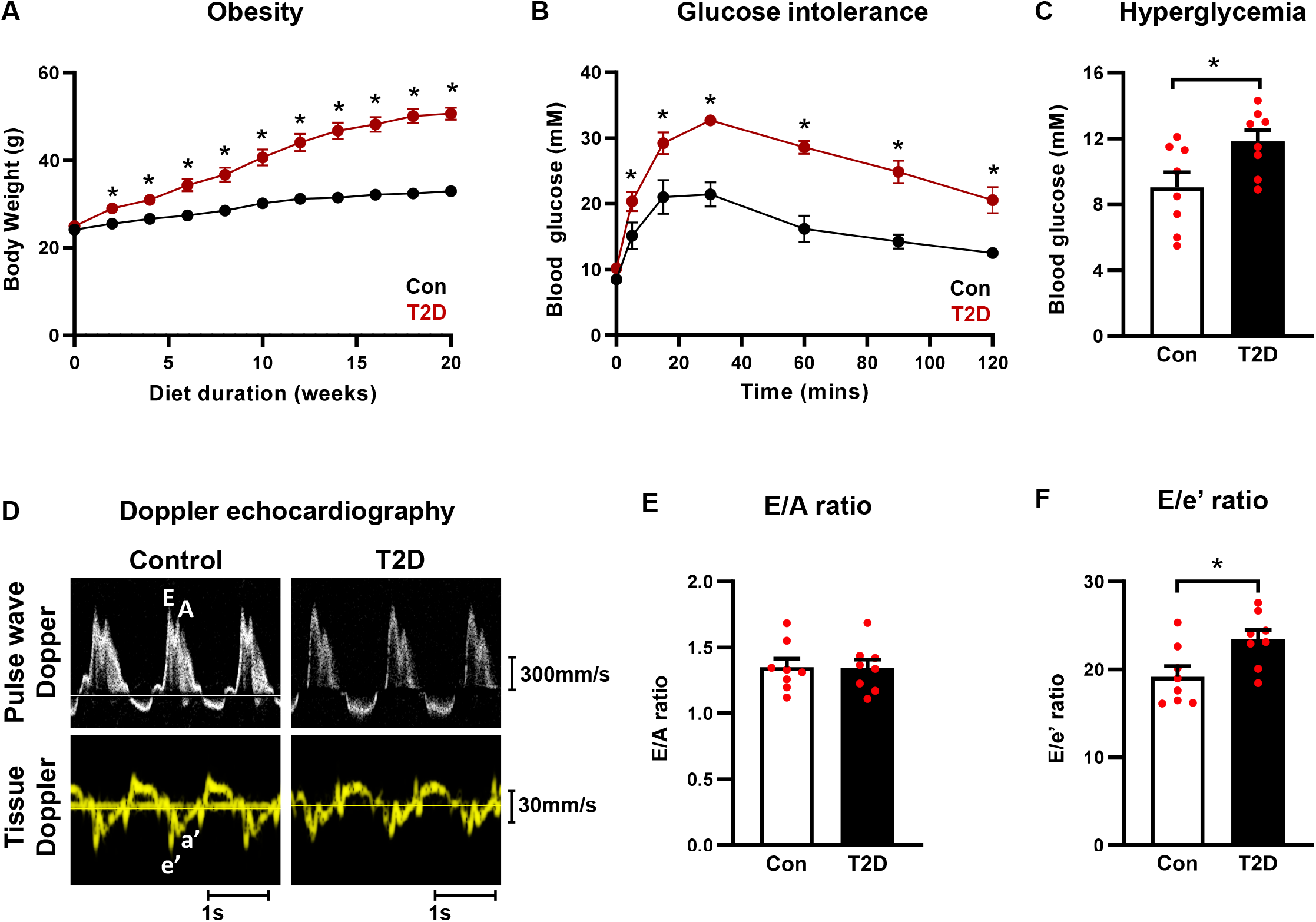
High fat high sugar diet-induced T2D mice exhibit obesity, impaired glucose tolerance, hyperglycemia and diastolic function. (**A**) Body weight progression throughout the 20 week dietary intervention. Note, in some instances error bars are not discernible as they fall within symbol shapes. Analyzed by 1-way repeated measures ANOVA, annotated with LSD *post hoc* analyses. (**B**) Glucose tolerance test in T2D mice (11 weeks high fat high sugar diet). Note, in some instances error bars are not discernible as they fall within symbol shapes. Analyzed by 1-way repeated measures ANOVA, annotated with LSD *post hoc* analyses. (**C**) Blood glucose levels in T2D mice at study endpoint (20 weeks high fat high sugar diet). Analyzed by Students unpaired t-test. (**D**) Representative Doppler echocardiography traces for control and T2D mice. Upper panels, mitral valve pulse wave (blood flow) Doppler imaging. Lower panels, mitral annulus tissue Doppler imaging. (**E**) Ratio of pulse wave Doppler E wave to A wave amplitude in T2D mice (20 weeks high fat high sugar diet). Analyzed by Students unpaired t-test. (**F**) Ratio of mitral valve flow Doppler E wave to mitral annulus tissue Doppler e’ wave in T2D mice (20 weeks high fat high sugar diet). Analyzed by Students unpaired t-test. Data are presented as mean ± SEM. n = 8/group. *p< 0.05.

### T2D mice exhibit preserved ejection fraction and reduced global longitudinal strain

To evaluate cardiac dimensions and systolic function, we performed M-mode echocardiography and B-mode 2D speckle tracking strain imaging in control and T2D mice. Exemplar M-mode and B-mode echocardiography traces are shown in Fig. 2A. Left ventricular hypertrophy was detected in T2D mice, as evidenced by increased left ventricular mass, and increased systolic and diastolic posterior wall thickness (Table 1). Systolic function measured via ejection fraction (Fig 2B) and fractional shortening (Table 1) was not different between control and T2D mice. Global strain measured in the short-axis view (radial and circumferential planes) was not different between control and T2D mice (Figs. 2C, 2D, Supp Table 1). In contrast, global longitudinal strain (Fig. 2E, Supp Table 1) measured in the long-axis view was significantly reduced in the T2D mice (T2D: -21.4 ± 1.1 % vs Control: -25.6 ± 1.2 %, p<0.05). These data suggest that T2D mice exhibit some evidence of mild longitudinal systolic dysfunction, despite preserved ejection fraction.

**Table 1.** In vivo echocardiography cardiac functional and structural characteristics in T2D mice. ESV, left ventricular end systolic volume; EDV, left ventricular end diastolic volume; LVAWs, left ventricular anterior wall thickness at end systole; LVAWd, left ventricular anterior wall thickness at end diastole; LVPWs, left ventricular posterior wall thickness at end systole; LVPWd, left ventricular posterior wall thicknesss at end diastole; E wave, mitral valve blood flow velocity during the early ventricular filling phase; A wave, mitral valve blood flow velocity during atrial contraction; e’ wave, mitral annulus tissue movement velocity during the early ventricular filling phase; a’ wave, mitral annulus tissue movement velocity during atrial contraction. Data are presented as mean ± SEM. n = 8/group. Analyzed by Students unpaired t-test, *p< 0.05.

|  | Control | T2D |
| --- | --- | --- |
| Heart rate (bpm) | 508 ± 8.5 | 486 ± 6.7 |
| LV mass (corrected) (mg) | 81.6 ± 25 | 103.6 ± 15 * |
| ESV (μl) | 16.3 ± 3.2 | 12.0 ± 1.3 |
| EDV (μl) | 54.1 ± 6.6 | 50.2 ± 1.9 |
| Fractional shortening (%) | 41.8 ± 2.5 | 45.4 ± 2.3 |
| Stroke volume (μl) | 37.8 ± 3.6 | 38.2 ± 1.1 |
| Cardiac output (ml/min) | 19.3 ± 2.0 | 18.5 ± 0.6 |
| LVAWs | 1.44 ± 0.059 | 1.66 ± 0.087 |
| LVAWd | 0.876 ± 0.040 | 1.04 ± 0.049 |
| LVPWs | 1.20 ± 0.046 | 1.42 ± 0.050 * |
| LVPWd | 0.712 ± 0.022 | 0.974 ± 0.038 * |
| E wave (mm/s) | 628 ± 28 | 705 ± 29 |
| A wave (mm/s) | 477 ± 39 | 537 ± 37 |
| e' wave (mm/s) | 33.5 ± 2.2 | 30.4 ± 1.6 |
| a' wave (mm/s) | 24.1 ± 2.1 | 20.7 ± 1.5 |

**Figure 2.**
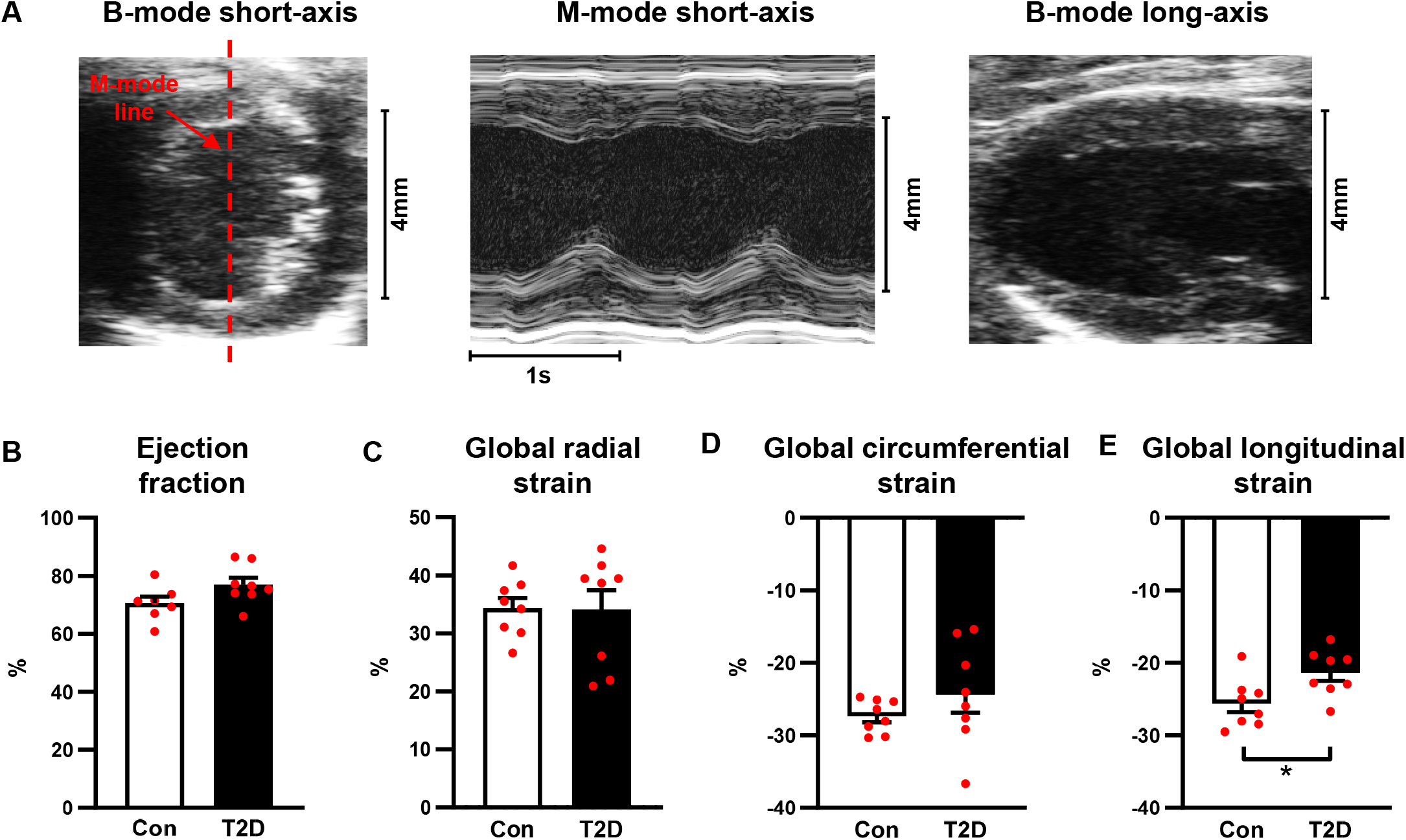
T2D mice exhibit longitudinal systolic dysfunction. (**A**) Exemplar M-mode and B-mode traces from the left ventricular short- and long-axis imaging planes. (**B**) Ejection fraction. (**C**) Global radial strain, analyzed in the short-axis imaging plane. (**D**) Global circumferential strain, analyzed in the short-axis imaging plane. (**E**) Global longitudinal strain, analyzed in the long-axis imaging plane. Data are presented as mean ± SEM. n = 7-8/group. Analyzed by Students unpaired t-test, *p< 0.05.

### Peak diastolic strain rate and peak diastolic velocity are impaired in T2D mice

A key advantage of speckle tracking echocardiography is the ability to measure longitudinal, radial and circumferential myocardial deformation throughout the cardiac cycle, and therefore distinct measures of systolic and diastolic ventricular kinetics can be acquired. Longitudinal peak diastolic strain rate and velocity were reduced in T2D mice (33% and 32% decrease respectively, p<0.05, Figs. 3A, 3C, Supp Table 1). Longitudinal peak systolic strain rate was unchanged (Fig. 3B, Supp Table 1) and a small but significant decrease in peak systolic velocity was detected (16% decrease, p<0.05, Fig. 3D, Supp Table 1). A schematic of regional segmentation of the ventricle in long axis view is shown in Fig. 3E. Regional analysis revealed that changes in longitudinal peak diastolic strain rate were most prominent in the anterior free wall (apex and mid regions, Fig. 3F).

**Figure 3.**
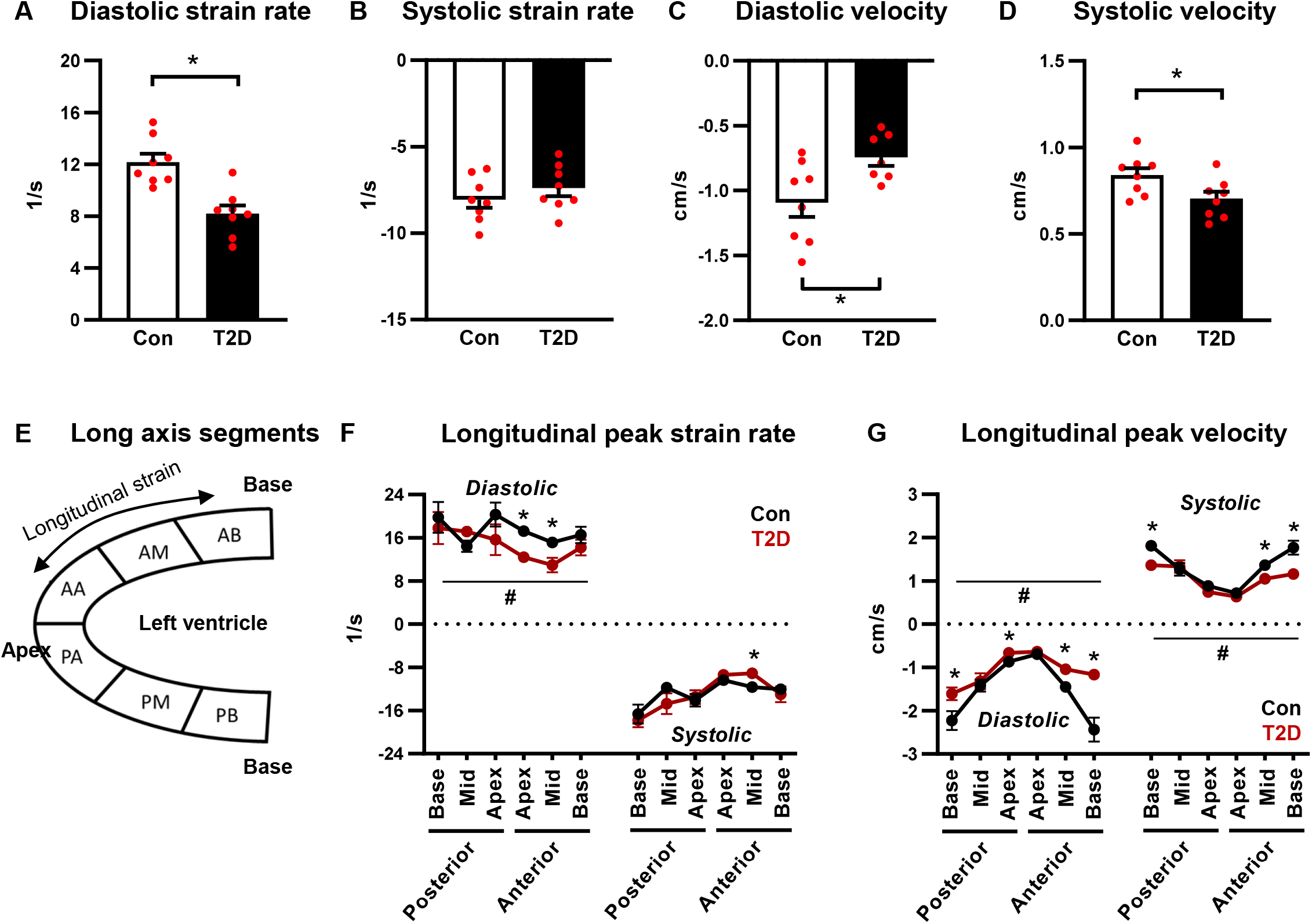
Longitudinal peak diastolic strain rate and velocity are reduced in T2D mice. (**A**) Exemplar longitudinal strain rate and velocity curves. (**B**) Schematic of long axis view left ventricle segmentation for strain analysis. (**C-D**) Longitudinal peak diastolic and systolic strain rate. Analyzed by Students unpaired t-test. (**E-F**) Longitudinal peak diastolic and systolic velocity. Analyzed by Students unpaired t-test. (**G**) Regional longitudinal peak strain rate. Analyzed by 1-way repeated measures ANOVA, annotated with LSD *post hoc* analyses. (**H**) Regional longitudinal peak velocity. Analyzed by 1-way repeated measures ANOVA, annotated with LSD *post hoc* analyses. Data are presented as mean ± SEM. n = 7-8/group. *p<0.05 *post hoc* analysis; ^#^p<0.05 T2D factor effect. PB: posterior base; PM: posterior middle; PA: posterior apex; AA: anterior apex; AM: anterior middle; AB: anterior base.

Longitudinal peak diastolic velocity was significantly reduced in the posterior base and apex, and anterior mid- and base regions (Fig. 3G). Reduced peak systolic velocity was detected in similar regions (Fig. 3G). An overall significant T2D ANOVA factor effect was evident for reduced longitudinal peak diastolic strain rate, peak diastolic velocity and peak systolic velocity (Figs. 3F-G), consistent with global measures presented in Figs. 3C-D.

Radial diastolic peak strain rate and velocity were reduced in T2D mice (33% and 26% decrease respectively, p<0.05, Figs. 4A, 4C, Supp Table 1). Radial peak systolic strain rate and velocity were unchanged (Figs. 4B, 4D, Supp Table 1). A schematic of regional segmentation of the ventricle in long axis view is shown in Fig. 4E. Regional analysis of radial peak diastolic and systolic strain rate detected no significant differences between control and T2D mice (Fig. 4F). An overall significant T2D ANOVA factor effect was evident for peak diastolic velocity, and this effect was most marked in the anterior apex region (Fig. 4G).

**Figure 4.**
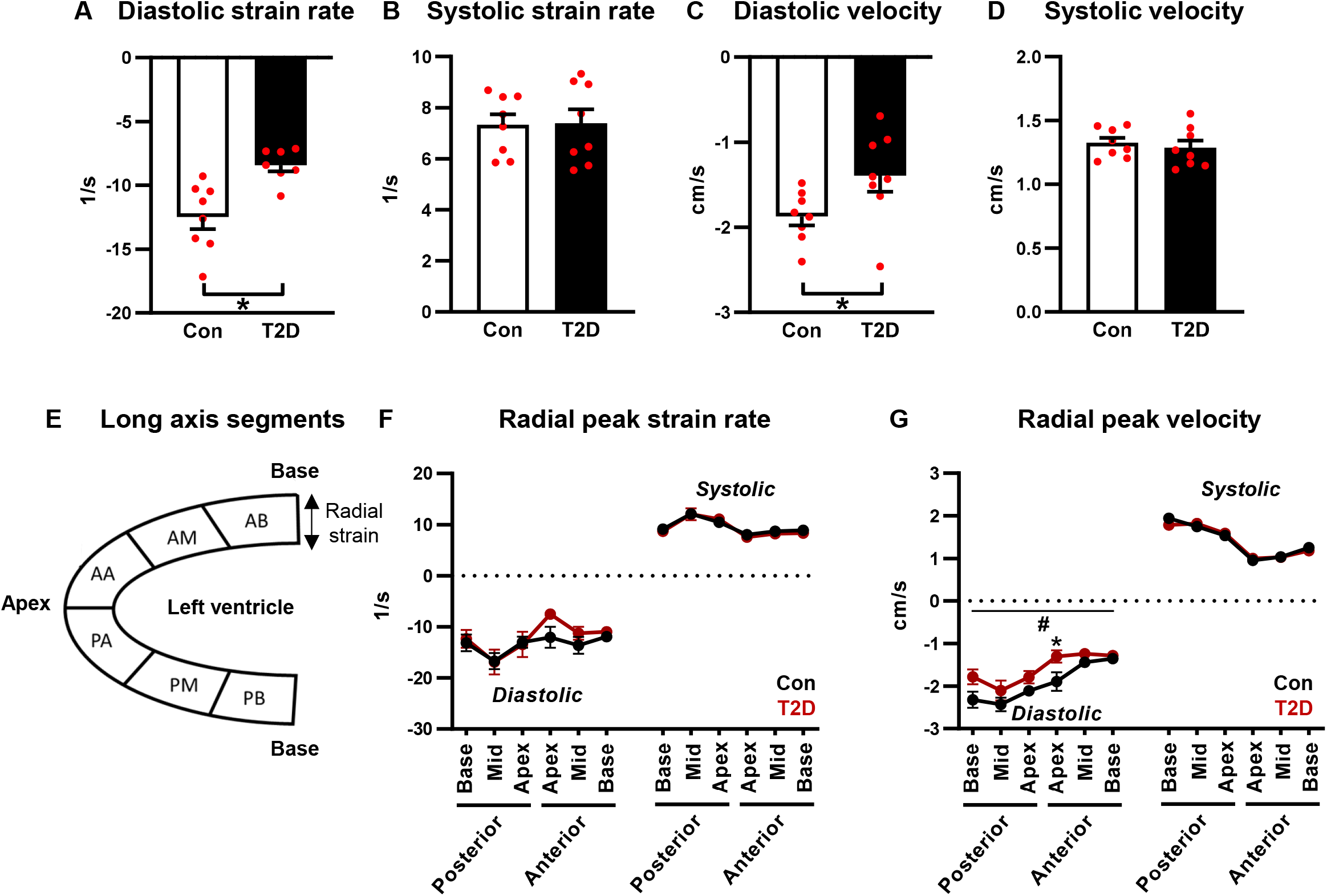
Radial diastolic peak strain rate and velocity are reduced in T2D mice. (**A**) Exemplar radial strain rate and velocity curves. (**B**) Schematic of long axis view left ventricle segmentation for strain analysis. (**C-D**) Radial diastolic and systolic peak strain rate. Analyzed by Students unpaired t-test. (**E-F**) Radial diastolic and systolic peak velocity. Analyzed by Students unpaired t-test. (**G**) Regional radial peak strain rate. Analyzed by 1-way repeated measures ANOVA, annotated with LSD *post hoc* analyses. (**H**) Regional radial peak velocity. Analyzed by 1-way repeated measures ANOVA, annotated with LSD *post hoc* analyses. Data are presented as mean ± SEM. n = 7-8/group. *p<0.05 *post hoc* analysis; ^#^p<0.05 T2D factor effect. PB: posterior base; PM: posterior middle; PA: posterior apex; AA: anterior apex; AM: anterior middle; AB: anterior base.

Circumferential diastolic peak strain rate was reduced in T2D mice (34% decrease, p<0.05, Fig. 5A, Supp Table 1) and diastolic peak velocity was not different between groups (Fig. 5C, Supp Table 1). Circumferential peak systolic strain rate and velocity were unchanged (Figs. 5B, 5D, Supp Table 1). A schematic of regional segmentation of the ventricle in short axis view is shown in Fig. 5E. Regional analysis of circumferential diastolic peak strain rate detected significantly lower values in the inferior free wall and anterior septum regions (Fig. 5F) and diastolic peak velocity was significantly lower in the posterior septal wall (Fig. 5G). A small but significant decrease in circumferential systolic peak velocity was evident in the posterior septal wall region (Fig. 5G). Collectively these data on myocardial wall deformation kinetics confirm that diastolic dysfunction is detected using peak strain rate and velocity analyses.

**Figure 5.**
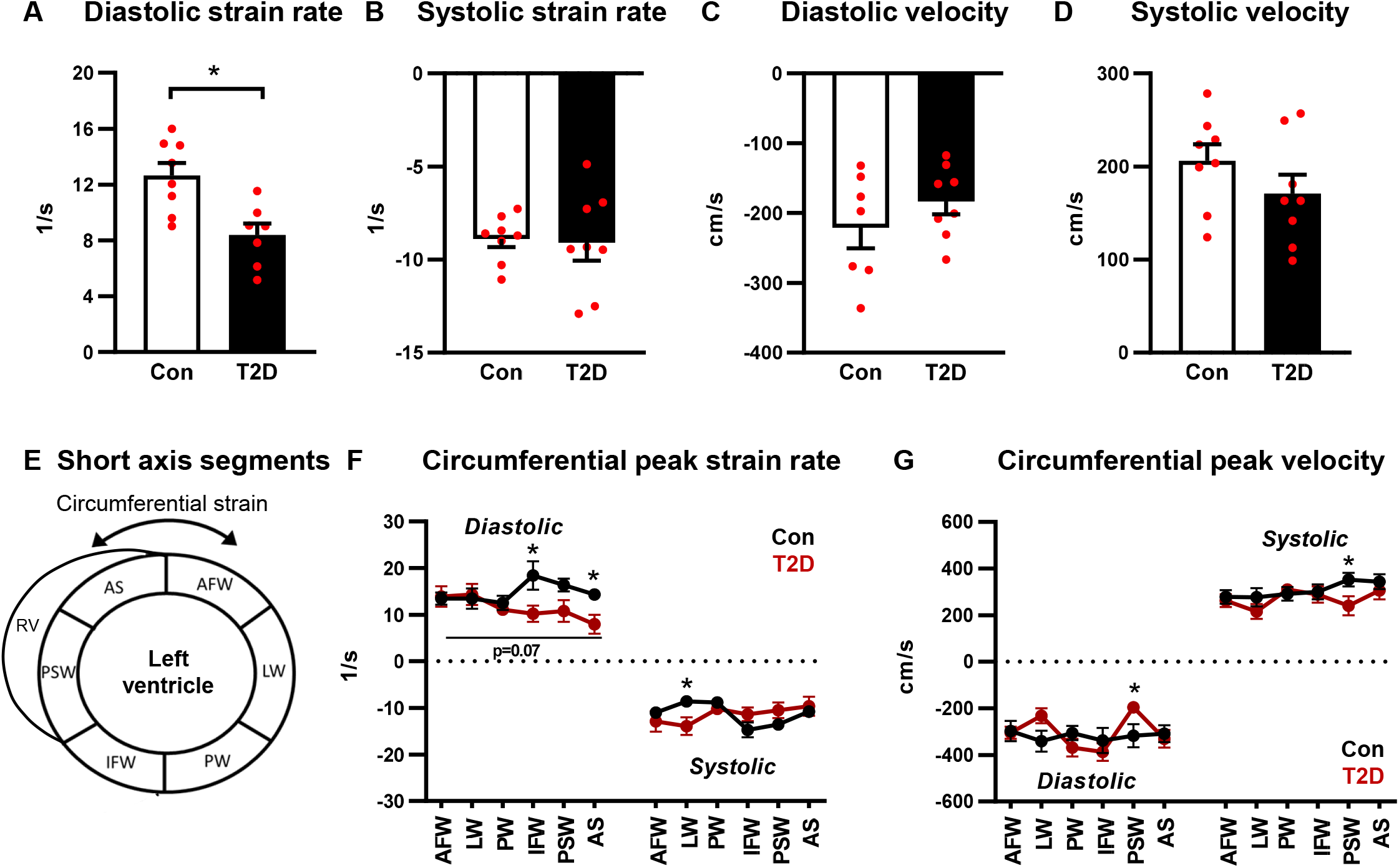
Circumferential diastolic strain rate and velocity are reduced in T2D mice. (**A**) Exemplar circumferential strain rate and velocity curves. (**B**) Schematic of short axis view left ventricle segmentation for strain analysis. (**C-D**) Circumferential diastolic and systolic peak strain rate. Analyzed by Students unpaired t-test. (**E-F**) Circumferential diastolic and systolic peak velocity. Analyzed by Students unpaired t-test. (**G**) Regional circumferential peak strain rate. Analyzed by 1-way repeated measures ANOVA, annotated with LSD *post hoc* analyses. P=0.07 T2D factor effect. (**H**) Regional circumferential peak velocity. Analyzed by 1-way repeated measures ANOVA, annotated with LSD *post hoc* analyses. Data are presented as mean ± SEM. n = 7-8/group. *p< 0.05. RV: right ventricle; AFW: anterior free wall; LW: lateral wall; PW: posterior wall; IFW: inferior free wall; PSW: posterior septal wall; AS: anterior septum.

### Speckle tracking-derived measures of diastolic dysfunction are correlated with E/e’

To determine whether measures of diastolic function using speckle tracking echocardiography are comparable to tissue Doppler imaging methods, we investigated the relationship between E/e’ and diastolic peak strain rate and velocity. Significant correlations between E/e’ and longitudinal diastolic peak strain rate and velocity were evident (Figs. 6A, 6B), suggesting that speckle tracking echocardiography may be a suitable surrogate for Doppler imaging for assessment of diastolic function in diabetic mice.

**Figure 6.**
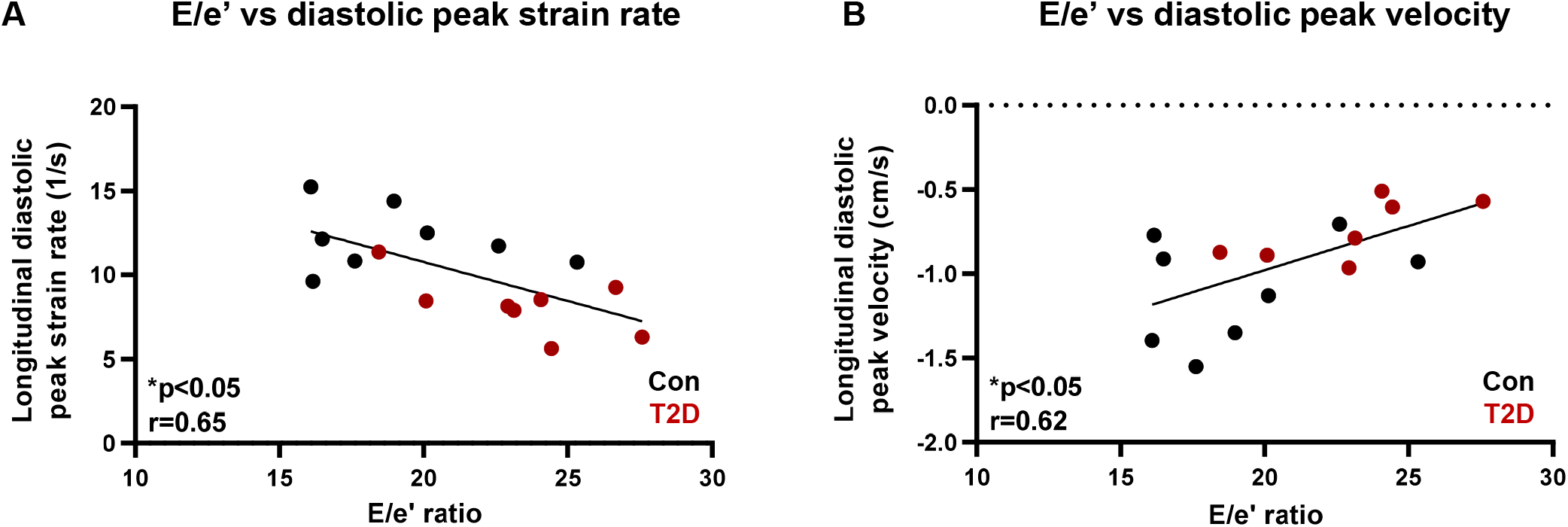
Speckle tracking echocardiography is comparable to doppler measures of diastolic function. (**A**) E/e’ and longitudinal diastolic peak strain rate correlation. (**B**) E/e’ and longitudinal diastolic peak velocity correlation. n = 7-8/group. *p<0.05. r, Pearson correlation coefficient.

## Discussion

This study provides a comprehensive report of global and regional diastolic dysfunction in murine cardiopathology using 2D speckle tracking echocardiography. Peak diastolic strain rate was lower in T2D mice, evident in the longitudinal, radial and circumferential planes. Peak diastolic velocity was lower in T2D mice in the longitudinal and radial, but not circumferential planes. Consistent with the finding that ejection fraction was preserved in this setting, systolic strain rate was unchanged with T2D in all imaging planes. Interestingly, global longitudinal strain, typically considered a measure of systolic function, was significantly impaired in T2D mice, which may provide an indicator of early systolic impairment in the longitudinal plane. Regional analysis of longitudinal strain rate revealed that the anterior free wall of the left ventricle is particularly vulnerable to T2D-induced diastolic dysfunction. Longitudinal diastolic strain rate and velocity were closely correlated with the commonly reported Doppler index of diastolic dysfunction, E/e’. These findings deliver important benchmark values for future pre-clinical cardiology studies in the field of diastolic dysfunction, HFpEF and diabetic heart disease.

Although speckle tracking echocardiography was first introduced as a tool for *in vivo* assessment of myocardial strain in mice over 15 years ago (Li et al 2007, Peng et al 2009), very few studies have used this technique to report specific measures of diastolic function in rodents, i.e. diastolic peak strain rate and diastolic peak velocity. In the present study, longitudinal diastolic peak strain rate and velocity significantly correlated with the commonly reported index of diastolic dysfunction derived from Doppler imaging, E/e’. Overall, the extent of reduction in diastolic strain rate (longitudinal, radial & circumferential, 33-34% decrease) was larger than the extent of increase in E/e’ detected in diabetic mice (22% increase). The use of high frequency echocardiography for measurement of diastolic strain rate and velocity provides an angle-independent method of assessing diastolic dysfunction in settings of fast heart rates in mice (>400 bpm). The pulse-wave Doppler signal is influenced by alignment of the ultrasound beam with the flow of blood and is heart rate dependent. Further, E/e’ is measured near the mitral annulus and consequently does not capture the global relaxation properties of the heart^18^. The acquisition of high quality b-mode images for strain analysis is largely dependent on the resolution and frame rates of the echocardiography platform, and is less dependent on the operator than Doppler methods. Post-processing of b-mode images does however incorporate some element of operator subjectivity, and development of artificial intelligence tools for strain analysis has emerged in the clinical literature to overcome this limitation^19,20^.

In the pre-clinical literature, regional analysis of myocardial wall deformation has primarily been used in studies of myocardial infarction for identifying areas of dyssynchrony in the ventricular wall.^11^ In the present study, regional analysis revealed that impairment in longitudinal diastolic strain rate in diabetic mice was most prominent in the anterior free wall. Impairment in circumferential diastolic strain rate was most prominent in the inferior free wall and anterior septum. These findings are similar to regional changes in myocardial deformation reported in adolescents with type 1 diabetes using cardiac MRI.^21^ Significant impairment in diastolic strain rate was observed in the inferior septal and free wall regions.^21^ This previous study calculated the diastolic relaxation fraction (ratio of myocardial contraction to relaxation during the filling phase^22^) and reported that adolescent T1D patients exhibit early diastolic left ventricular discoordination, despite otherwise normal cardiac function and left ventricular size. Thus assessment of regional myocardial wall deformation may provide an early indicator of diastolic dysfunction by identifying disparate regions of impairment.

While left ventricular ejection fraction has been the most widely used clinical measure of cardiac function for decades, global longitudinal strain is increasingly recognized as an important clinical indicator, with prognostic value in predicting adverse outcomes in chronic systolic heart failure^23,24^. Global longitudinal strain quantifies the maximum change in length of the left ventricle during contraction (as a percentage of the initial LV dimension), thus is typically considered a measure of systolic function. Interestingly, in heart failure patients where ejection fraction is preserved (HFpEF) and systolic function is generally considered to be normal, global longitudinal strain is frequently reported to be decreased. This could reflect a difference in the imaging planes used.

Ejection fraction is typically measured using M-mode analysis in a short-axis view, while global longitudinal strain is measured in the long-axis view. In the present study, measures of systolic function acquired from the short-axis imaging plane (ejection fraction, global circumferential strain, global radial strain) were unchanged in diabetic mice. In contrast, global longitudinal strain was reduced, suggesting that longitudinal systolic dysfunction is present. These findings are consistent with other pre-clinical literature reporting that diabetic mice with normal ejection fraction exhibit reduced global longitudinal strain and diastolic dysfunction^25,26^. Clinically, global longitudinal strain has been incorporated into recently updated European Society of Cardiology guidelines for diagnosis of HFpEF^27^. In chronic settings of progressive cardiac dysfunction, such as diabetes, global longitudinal strain may provide a robust measure of early subclinical systolic dysfunction, with high predictive value.

In the present study, both peak diastolic strain rate and peak diastolic velocity have been incorporated into the assessment of diastolic dysfunction in diabetic mice. In most cases, these variables show the same impairment, with the exception of unchanged circumferential peak diastolic velocity. By definition, strain rate is the derivative of strain, and is therefore presented as relative to the initial dimension (L_0_). Given that velocity is the rate of change of wall motion, it is therefore not dependent on the initial size of the ventricular segment. Thus, in a setting of diabetes where changes in the ventricular wall dimensions would be expected (as is the case in the present study), subtle differences in strain rate and velocity may be detectable. Additionally, there is some evidence from the clinical literature that strain rate may be more independent of acute changes in loading conditions (preload & afterload) than tissue velocities^28^, but further work is required to validate these findings in chronic settings.

## Conclusion

In conclusion, here we provide evidence to support the utility of 2D speckle tracking echocardiography to document global and regional diastolic dysfunction in a pre-clinical setting of murine cardiopathology. Peak diastolic strain rate and peak diastolic velocity constitute reliable indicators of diastolic dysfunction in diabetic mice, significantly correlated with traditional Doppler measures of impaired ventricular relaxation. Some evidence of longitudinal systolic impairment was detected by global longitudinal strain, despite preserved ejection fraction. The anterior free left ventricular wall appeared to be most vulnerable to longitudinal diastolic impairment. These findings provide a significant advance on characterization of diastolic dysfunction in a pre-clinical mouse model of diabetes and offer a comprehensive suite of benchmark values for future pre-clinical cardiology studies.

## Methods

### Animals

All animal experiments were approved by the University of Auckland Animal Ethics Committee and complied with the guidelines and regulations of the Code of Practice for the Care and Use of Animals for Scientific Purposes. Animals were randomly assigned to experimental groups and were group housed (4 mice per cage) in a temperature-controlled environment with 12-hour light/dark cycles. Food and water was provided *ad libitum*.

#### Type 2 diabetic mice

Type 2 diabetes was induced in C57Bl/6J mice by a high fat high sugar dietary intervention commencing at 7 weeks of age. After an initial 1 week transitional feeding period, animals were fed a high fat high sugar diet (43% kcal from fat, 200g/kg sucrose, SF04-001, Specialty Feeds, Australia – based on the formulation of Research Diets D12451, USA) or control reference diet (16% kcal from fat, 100g/kg sucrose, custom mouse AIN93G control diet, Specialty Feeds, Australia) for 20 weeks duration. Fresh diet was provided twice a week.

### Glucose tolerance testing

Glucose tolerance testing was performed after 11 weeks of dietary intervention following 6 hours fasting. Baseline blood glucose levels were measured using an Accu-Check glucometer with a small blood sample from a needle prick to the tail vein. Glucose (1.5 g/kg body weight) was injected i.p. and blood glucose was measured at 5, 10, 30, 60, 90 and 120 minutes after the glucose injection.

### Echocardiography

Animals were anaesthetized with isoflurane and transthoracic echocardiography was performed using the VEVO LAZR-X 3100 with a MX400 probe (20-46 Hz) linear array transducer coupled with digital ultrasound system (FUJIFLIM Visual Sonics). All analyses were performed by an investigator blinded to study group allocation.

#### M-mode echocardiography imaging

Left ventricular (LV) M-mode two-dimensional echocardiography was performed in a parasternal short axis view at the mid papillary level to measure LV wall and chamber dimensions to derive systolic function parameters: % ejection fraction ((end diastolic volume-end systolic volume)/end diastolic volume) x 100), and % fractional shortening ((LV end diastolic diameter-LV end systolic diameter)/(LV end diastolic diameter) x100). LV mass was calculated: 1.053 x (LV end diastolic diameter + LV end diastolic posterior wall thickness + end diastolic inter-ventricular septal wall thickness)^3^ – end diastolic diameter^3^, with correction factor 0.8. At least 3 consecutive cardiac cycles were sampled per cineloop image and 3 images per animal were analyzed and averaged.

#### Pulse wave Doppler and tissue Doppler imaging

Pulse wave Doppler and tissue Doppler imaging were acquired from the apical 4 chamber view to assess LV diastolic function parameters: velocity of mitral inflow during early passive filling (E) and during atrial contraction (A), and velocity of mitral annulus during early passive filling (e’) and during atrial contraction (a’). At least 3 cardiac cycles were sampled per cineloop image and 2-4 images per animal were analyzed and averaged. Wave forms from trans-mitral and tissue records were evaluated with temporal registration to validate waveform identification.

#### Speckle tracking echocardiography

Two-dimensional echocardiography B-mode loops were acquired from the left ventricular long axis (longitudinal and radial strain) and short axis (circumferential strain) views, and analyzed using Vevo strain software (VisualSonics). Endocardial and epicardial boarders were traced and checked throughout three cardiac cycles to ensure optimal tracking. Strain and displacement analyses were performed providing global and segmental (regional) values. Strain rate and velocity parameters were derived from the strain-time and displacement-time relations respectively. Average values were calculated from 2-3 independent images per animal.

### Statistics

Data are presented as mean ± SEM and statistical analysis was performed using Graphpad Prism v7.0. All data sets were tested for normal distribution using Shapiro-Wilk test. Differences between two groups were analyzed by Students unpaired t-test. For analysis of body weight, glucose tolerance and regional strain parameters, a repeated measures ANOVA with LSD *post hoc* tests was used. For correlation analyses, Pearson’s correlation coefficient was used. A p-value of <0.05 was considered statistically significant.

## Acknowledgements

The research was supported by funding from the Health Research Council of New Zealand (19/190) and Marsden (19-UOA-268). The authors acknowledge Justin Menezes who contributed to initial echocardiographic image analysis.

## Author contributions

L.J.D., C.M., K.M.M. conceived and designed the research. L.J.D. and C.M. performed animal monitoring, echocardiography scans and image analysis. L.J.D., C.M., P.K., M.A., A.J.A.R., K.L.W., J.R.B., J.V.J., C.L.C., L.M.D.D., K.M.M. interpreted results of experiments. L.J.D. and K.M.M. drafted the manuscript. All authors approved the final version.

## Data availability statement

The datasets generated and analyzed during the current study are available from the corresponding author on reasonable request.

## References

1. Jackson, A. M., Rorth, R., Liu, J., Kristensen, S. L., Anand, I. S., Claggett, B. L., Cleland, J. G. F., Chopra, V. K., Desai, A. S., Ge, J., Gong, J., Lam, C. S. P., Lefkowitz, M. P., Maggioni, A. P., Martinez, F., Packer, M., Pfeffer, M. A., Pieske, B., Redfield, M. M., Rizkala, A. R., Rouleau, J. L., Seferovic, P. M., Tromp, J., Van Veldhuisen, D. J., Yilmaz, M. B., Zannad, F., Zile, M. R., Kober, L., Petrie, M. C., Jhund, P. S., Solomon, S. D., McMurray, J. J. V., Committees, P.-H. & Investigators. Diabetes and pre-diabetes in patients with heart failure and preserved ejection fraction. Eur J Heart Fail. 24, 497–509, (2022).

2. McHugh, K., DeVore, A. D., Wu, J., Matsouaka, R. A., Fonarow, G. C., Heidenreich, P. A., Yancy, C. W., Green, J. B., Altman, N. & Hernandez, A. F. Heart Failure With Preserved Ejection Fraction and Diabetes: JACC State-of-the-Art Review. J Am Coll Cardiol. 73, 602–611, (2019).

3. Chandramouli, C., Reichelt, M. E., Curl, C. L., Varma, U., Bienvenu, L. A., Koutsifeli, P., Raaijmakers, A. J. A., De Blasio, M. J., Qin, C. X., Jenkins, A. J., Ritchie, R. H., Mellor, K. M. & Delbridge, L. M. D. Diastolic dysfunction is more apparent in STZ-induced diabetic female mice, despite less pronounced hyperglycemia. Sci Rep. 8, 2346, (2018).

4. De Blasio, M. J., Huynh, N., Deo, M., Dubrana, L. E., Walsh, J., Willis, A., Prakoso, D., Kiriazis, H., Donner, D. G., Chatham, J. C. & Ritchie, R. H. Defining the Progression of Diabetic Cardiomyopathy in a Mouse Model of Type 1 Diabetes. Frontiers in physiology. 11, 124, (2020).

5. Schnelle, M., Catibog, N., Zhang, M., Nabeebaccus, A. A., Anderson, G., Richards, D. A., Sawyer, G., Zhang, X., Toischer, K., Hasenfuss, G., Monaghan, M. J. & Shah, A. M. Echocardiographic evaluation of diastolic function in mouse models of heart disease. Journal of molecular and cellular cardiology. 114, 20–28, (2018).

6. Mondillo, S., Galderisi, M., Mele, D., Cameli, M., Lomoriello, V. S., Zacà, V., Ballo, P., D’Andrea, A., Muraru, D., Losi, M., Agricola, E., D’Errico, A., Buralli, S., Sciomer, S., Nistri, S. & Badano, L. Speckle-tracking echocardiography: a new technique for assessing myocardial function. Journal of ultrasound in medicine : official journal of the American Institute of Ultrasound in Medicine. 30, 71–83, (2011).

7. Lorenzo-Almorós, A., Tuñón, J., Orejas, M., Cortés, M., Egido, J. & Lorenzo, Ó. Diagnostic approaches for diabetic cardiomyopathy. Cardiovascular diabetology. 16, 28, (2017).

8. Buggey, J., Alenezi, F., Yoon, H. J., Phelan, M., DeVore, A. D., Khouri, M. G., Schulte, P. J. & Velazquez, E. J. Left ventricular global longitudinal strain in patients with heart failure with preserved ejection fraction: outcomes following an acute heart failure hospitalization. ESC heart failure. 4, 432–439, (2017).

9. Moharram, M. A., Lamberts, R. R., Whalley, G., Williams, M. J. A. & Coffey, S. Myocardial tissue characterisation using echocardiographic deformation imaging. Cardiovasc Ultrasound. 17, 27, (2019).

10. Peng, Y., Popovic, Z. B., Sopko, N., Drinko, J., Zhang, Z., Thomas, J. D. & Penn, M. S. Speckle tracking echocardiography in the assessment of mouse models of cardiac dysfunction. Am J Physiol Heart Circ Physiol. 297, H811–820, (2009).

11. Popovic, Z. B., Benejam, C., Bian, J., Mal, N., Drinko, J., Lee, K., Forudi, F., Reeg, R., Greenberg, N. L., Thomas, J. D. & Penn, M. S. Speckle-tracking echocardiography correctly identifies segmental left ventricular dysfunction induced by scarring in a rat model of myocardial infarction. Am J Physiol Heart Circ Physiol. 292, H2809–2816, (2007).

12. Bauer, M., Cheng, S., Jain, M., Ngoy, S., Theodoropoulos, C., Trujillo, A., Lin, F. C. & Liao, R. Echocardiographic speckle-tracking based strain imaging for rapid cardiovascular phenotyping in mice. Circ Res. 108, 908–916, (2011).

13. Bhan, A., Sirker, A., Zhang, J., Protti, A., Catibog, N., Driver, W., Botnar, R., Monaghan, M. J. & Shah, A. M. High-frequency speckle tracking echocardiography in the assessment of left ventricular function and remodeling after murine myocardial infarction. Am J Physiol Heart Circ Physiol. 306, H1371–1383, (2014).

14. Dandel, M., Lehmkuhl, H., Knosalla, C., Suramelashvili, N. & Hetzer, R. Strain and strain rate imaging by echocardiography - basic concepts and clinical applicability. Current cardiology reviews. 5, 133–148, (2009).

15. Pappritz, K., Grune, J., Klein, O., Hegemann, N., Dong, F., El-Shafeey, M., Lin, J., Kuebler, W. M., Kintscher, U., Tschöpe, C. & Van Linthout, S. Speckle-tracking echocardiography combined with imaging mass spectrometry assesses region-dependent alterations. Scientific reports. 10, 3629, (2020).

16. Shepherd, D. L., Nichols, C. E., Croston, T. L., McLaughlin, S. L., Petrone, A. B., Lewis, S. E., Thapa, D., Long, D. M., Dick, G. M. & Hollander, J. M. Early detection of cardiac dysfunction in the type 1 diabetic heart using speckle-tracking based strain imaging. Journal of molecular and cellular cardiology. 90, 74–83, (2016).

17. Zhou, Y., Xiao, H., Wu, J., Zha, L., Zhou, M., Li, Q., Wang, M., Shi, S., Li, Y., Lyu, L., Wang, Q., Tu, X. & Lu, Q. Type I Diabetic Akita Mouse Model is Characterized by Abnormal Cardiac Deformation During Early Stages of Diabetic Cardiomyopathy with Speckle-Tracking Based Strain Imaging. Cellular physiology and biochemistry : international journal of experimental cellular physiology, biochemistry, and pharmacology. 45, 1541–1550, (2018).

18. Lassen, M. C. H., Jensen, M. T., Biering-Sorensen, T., Mogelvang, R., Fritz-Hansen, T., Vilsboll, T., Rossing, P. & Jorgensen, P. G. Prognostic value of ratio of transmitral early filling velocity to early diastolic strain rate in patients with Type 2 diabetes. Eur Heart J Cardiovasc Imaging, (2019).

19. Leeson, P. & Fletcher, A. J. Let AI Take the Strain. JACC Cardiovasc Imaging. 14, 1929–1931, (2021).

20. Salte, I. M., Ostvik, A., Smistad, E., Melichova, D., Nguyen, T. M., Karlsen, S., Brunvand, H., Haugaa, K. H., Edvardsen, T., Lovstakken, L. & Grenne, B. Artificial Intelligence for Automatic Measurement of Left Ventricular Strain in Echocardiography. JACC Cardiovasc Imaging. 14, 1918–1928, (2021).

21. Schafer, M., Bjornstad, P., Frank, B. S., Baumgartner, A., Truong, U., Enge, D., von Alvensleben, J. C., Mitchell, M. B., Ivy, D. D., Barker, A. J., Reusch, J. E. B. & Nadeau, K. J. Frequency of Reduced Left Ventricular Contractile Efficiency and Discoordinated Myocardial Relaxation in Patients Aged 16 to 21 Years With Type 1 Diabetes Mellitus (from the Emerald Study). Am J Cardiol. 128, 45–53, (2020).

22. Frank, B. S., Schafer, M., Douwes, J. M., Ivy, D. D., Abman, S. H., Davidson, J. A., Burzlaff, S., Mitchell, M. B., Morgan, G. J., Browne, L. P., Barker, A. J., Truong, U. & von Alvensleben, J. C. Novel measures of left ventricular electromechanical discoordination predict clinical outcomes in children with pulmonary arterial hypertension. Am J Physiol Heart Circ Physiol. 318, H401–H412, (2020).

23. Abou, R., van der Bijl, P., Bax, J. J. & Delgado, V. Global longitudinal strain: clinical use and prognostic implications in contemporary practice. Heart. 106, 1438–1444, (2020).

24. Ashish, K., Faisaluddin, M., Bandyopadhyay, D., Hajra, A. & Herzog, E. Prognostic value of global longitudinal strain in heart failure subjects: A recent prototype. Int J Cardiol Heart Vasc. 22, 48–49, (2019).

25. Zhang, X., Kong, S., Wu, M., Niu, Y., Wang, K., Zhu, H. & Yuan, J. Impact high fat diet on myocardial strain in mice by 2D speckle tracking imaging. Obes Res Clin Pract. 15, 133–137, (2021).

26. Soliman, H., Nyamandi, V., Garcia-Patino, M., Varela, J. N., Bankar, G., Lin, G., Jia, Z. & MacLeod, K. M. Partial deletion of ROCK2 protects mice from high-fat diet-induced cardiac insulin resistance and contractile dysfunction. Am J Physiol Heart Circ Physiol. 309, H70–81, (2015).

27. Pieske, B., Tschope, C., de Boer, R. A., Fraser, A. G., Anker, S. D., Donal, E., Edelmann, F., Fu, M., Guazzi, M., Lam, C. S. P., Lancellotti, P., Melenovsky, V., Morris, D. A., Nagel, E., Pieske-Kraigher, E., Ponikowski, P., Solomon, S. D., Vasan, R. S., Rutten, F. H., Voors, A., Ruschitzka, F., Paulus, W. J., Seferovic, P. & Filippatos, G. How to diagnose heart failure with preserved ejection fraction: the HFA-PEFF diagnostic algorithm: a consensus recommendation from the Heart Failure Association (HFA) of the European Society of Cardiology (ESC). Eur J Heart Fail. 22, 391–412, (2020).

28. Friedberg, M. K. & Mertens, L. Tissue velocities, strain, and strain rate for echocardiographic assessment of ventricular function in congenital heart disease. Eur J Echocardiogr. 10, 585–593, (2009).

