## Supplementary Table 1 for "Myocardial deformation imaging by 2D speckle tracking echocardiography for assessment of diastolic dysfunction in murine cardiopathology"

| | Control | T2D |
|---|---|---|
| Global radial strain (%) | 34.4 ± 1.7 | 34.1 ± 3.4 |
| Global circumferential strain (%) | -27.4 ± 0.8 | -24.4 ± 2.5 |
| Global longitudinal strain (%) | -25.6 ± 1.2 | -21.4 ± 1.1* |
| *Longitudinal:* | | |
| Diastolic peak strain rate (1/s) | 12.2 ± 0.6 | 8.20 ± 0.6* |
| Systolic peak strain rate (1/s) | -8.05 ± 0.5 | -7.39 ± 0.5 |
| Diastolic peak velocity (cm/s) | -1.09 ± 0.1 | -0.74 ± 0.1* |
| Systolic peak velocity (cm/s) | 0.84 ± 0.04 | 0.70 ± 0.04* |
| *Radial:* | | |
| Diastolic peak strain rate (1/s) | -12.5 ± 0.9 | -8.41 ± 0.50* |
| Systolic peak strain rate (1/s) | 7.33 ± 0.4 | 7.39 ± 0.6 |
| Diastolic peak velocity (cm/s) | -1.87 ± 0.1 | -1.39 ± 0.2* |
| Systolic peak velocity (cm/s) | 1.33 ± 0.04 | 1.29 ± 0.1 |
| *Circumferential:* | | |
| Diastolic peak strain rate (1/s) | 12.6 ± 0.9 | 8.39 ± 0.8* |
| Systolic peak strain rate (1/s) | -8.88 ± 0.4 | -9.09 ± 1.0 |
| Diastolic peak velocity (cm/s) | -221 ± 29 | -184 ± 18 |
| Systolic peak velocity (cm/s) | 206 ± 18 | 171 ± 20 |

**Supplementary Table 1.** Benchmark values of echocardiographic myocardial wall deformation imaging in T2D mice. Data are presented as mean ± SEM. n = 7-8/group. Analyzed by Students unpaired t-test, $^*p < 0.05$.
